# Large extracellular vesicles can be characterised by multiplex labelling using imaging flow cytometry

**DOI:** 10.1101/2020.02.07.938779

**Authors:** Suzanne M Johnson, Antonia Banyard, Christopher Smith, Aleksandr Mironov, Martin G McCabe

**Affiliations:** Children’s Cancer Group, Division of Cancer Sciences, School of Medical Sciences, Faculty of Biology Medicine and Health, University of Manchester, Oglesby Cancer Research Building, Manchester Academic Health Science Centre, Manchester Cancer Research Centre, Manchester, UK; Flow Cytometry Core Facility, Cancer Research UK Manchester Institute, University of Manchester, Alderley Park, Macclesfield, UK; Electron Microscopy Core Facility, School of Biological Sciences, Faculty of Biology Medicine and Health, University of Manchester, Manchester, M13 9PT, UK; Manchester Children’s Brain Tumour Research Network

## Abstract

Extracellular vesicles (EVs) are heterogeneous in size (30nm-10µm), content (lipid, RNA, DNA, protein) and potential function(s). Many isolation techniques routinely discard the large EVs at the early stages of small EV or exosome isolation protocols. We describe here a standardised method to isolate large EVs and examine EV marker expression and diameter using imaging flow cytometry.

**Methods:** We describe step-wise isolation and characterisation of a subset of large EVs from the medulloblastoma cell line UW228-2 assessed by fluorescent light microscopy, transmission electron microscopy (TEM) and TRPS. Viability of parent cells was assessed by Annexin V exposure by flow cytometry. Imaging flow cytometry (Imagestream Mark II) identified EVs by direct fluorescent membrane labelling with Cell Mask Orange (CMO) in conjunction with EV markers. A stringent gating algorithm based on side scatter and fluorescence intensity was applied and expression of EV markers CD63, CD9 and LAMP 1 assessed.

**Results:** UW228-2 cells prolifically release EVs of up to 6 µm. We show that the Imagestream Mark II imaging flow cytometer allows robust and reproducible analysis of large EVs, including assessment of diameter. We also demonstrate a correlation between increasing EV size and co-expression of markers screened.

**Conclusions:** We have developed a labelling and stringent gating strategy which is able to explore EV marker expression (CD63, CD9 and LAMP1) on individual EVs within a widely heterogeneous population. Taken together data presented here strongly support the value of exploring large EVs in clinical samples for potential biomarkers, useful in diagnostic screening and disease monitoring.

## INTRODUCTION

The term extracellular vesicle (EV) refers to particles released from cells which are delimited by a lipid bilayer, but do not contain a nucleus (1). EVs are heterogeneous in biogenesis (2), size and content. They range in size from 50nm exosomes (3) to oncosomes up to 10µm (4), and contain cargo of all biomolecule categories (5). Attempts to categorise EVs primarily by surface marker expression have been confounded by the recognition that many of the markers previously considered to be subset- or derivation-specific, are actually present on multiple or even all classes of EVs (6). Delineation and characterisation of specific EV subsets is an essential goal to achieve a better understanding of EV biology (7), yet there are no techniques that accurately quantify EVs across the full EV size range, or combine quantification with the ability to screen for EV marker expression or EV content.

Small EVs are characteristically isolated by high-speed centrifugation at 100,000 × g (8), and include both EVs derived intracellularly from late endosomes and released by exocytosis (exosomes), and other small EVs not derived from endosomes (3). Multiple commercial solutions exist for the isolation of exosomes from a variety of biological fluids including tissue culture supernatant, plasma and urine. In contrast, there are no commercially available solutions for the isolation of large EVs. As a result, isolation methods vary, and knowledge of large EV content and function in biological samples is relatively lacking. Large EVs, defined as >200 nm by recent guidelines set out by the International Society of Extracellular Vesicles (1), include cancer cell-derived oncosomes, dead cell-derived apoptotic bodies and platelets, and are visible by light microscopy (9). In published literature, EVs larger than 1 µm have historically been assumed to be apoptotic bodies (10). However, we and others have demonstrated that viable cell cultures produce large EVs which do not have the ultrastructural features reminiscent of fragments of apoptotic cells (4).

We previously reported a population of large EVs released by leukaemic cells which were actin-rich and contained intact organelles (11). These large EVs could be internalised by normal stromal cells and induced a switch in the preferred metabolic pathway of the recipient cells (9). Additionally, we found that leukaemia-derived EVs expressed a surface marker indicative of their parent cell (CD19) and could be detected in the peripheral blood of murine models and patient bone marrow plasma (9). Taken together, our previous work and existing large EV literature suggest that large EVs, often discarded in techniques to isolate smaller EVs and exosomes, could be considered as extensive reservoirs of biomolecules useful to study EV biogenesis and function, and to identify clinically relevant biomarkers for disease detection and treatment monitoring.

In this proof of concept study, we set out to: 1) highlight the abundance of large EVs produced by cells derived from the malignant brain tumour medulloblastoma *in vitro*; 2) describe variations in the expression of established EV markers in the large EV population; 3) describe how the Imagestream (ISX) can address sample heterogeneity by facilitating high throughput, single event EV analyses. We describe a protocol to isolate intact large EVs without cell contamination, from cells growing in serum-free medium, using gravity flow filtration combined with low-speed centrifugation.

We report for the first time a characterisation of size distribution and EV marker expression in this heterogeneous EV population, undertaken in accordance with the most recent international consensus guidelines for EV research from the International Society of Extracellular Vesicles (1).

## MATERIALS AND METHODS

### Cell line

UW228-2 cells were kindly provided by Dr DTW Jones (DKFZ, Heidleberg). Cells were grown in DMEM with L-glutamine (Lonza, UK cat: R8758) supplemented with 10% FCS (SIGMA, cat: F9665) in Corning 225 cm^2^ Angled Neck Cell Culture Flask with Vent Cap (cat: 431082) at 37 °C with 5% CO_2_ in normoxia. Cell lines were passaged with 1 × Trypsin-EDTA (Lonza, UK cat: T3924). All cell lines tested negative for Mycoplasma and all were authenticated in-house (CRUK-Manchester Institute) by examination of a total of 21 loci across the genome using the Powerplex 21 System (Promega).

### Chemicals and reagents including antibodies

See Table 1

**TABLE 1:**
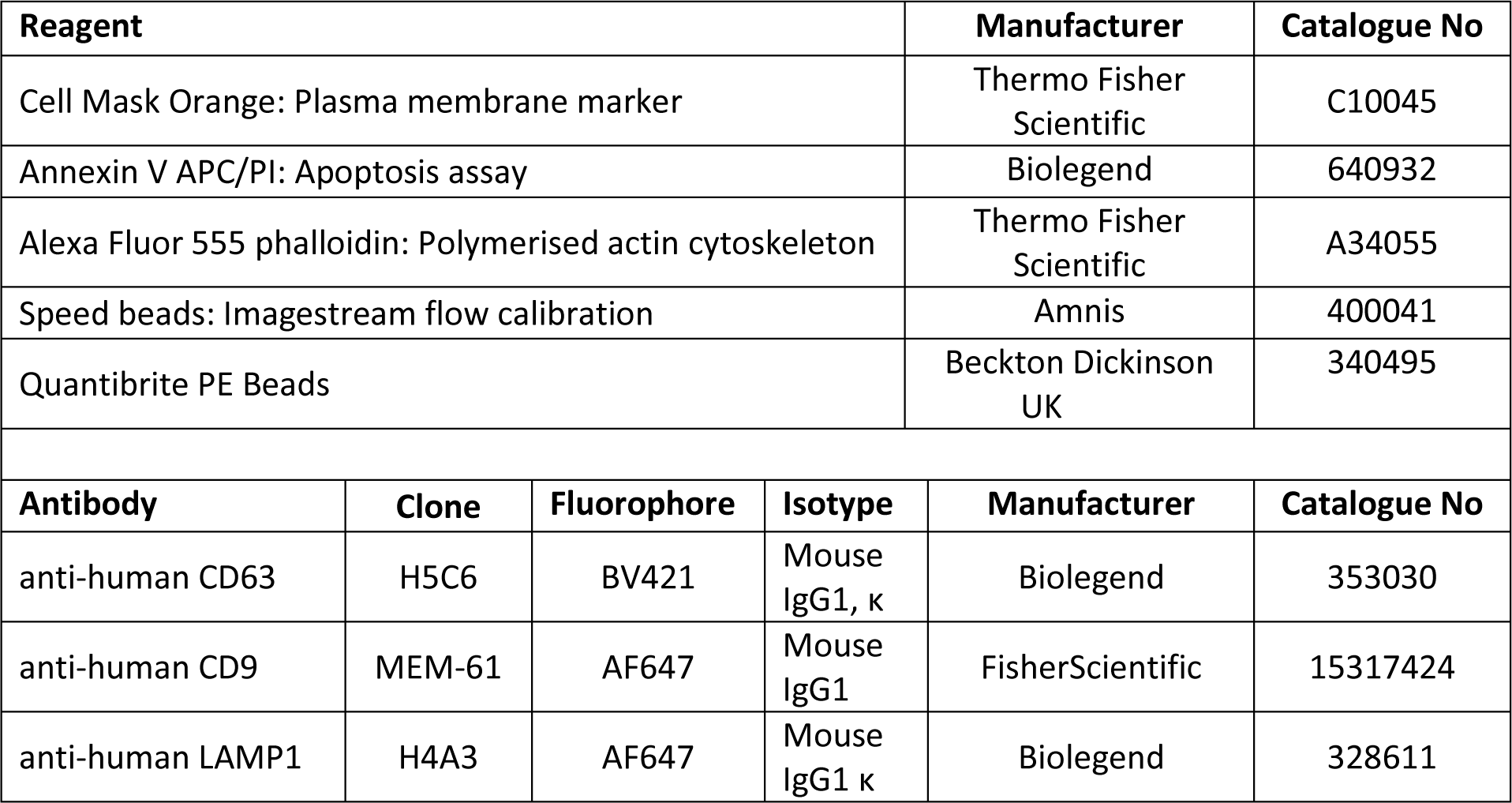
Reagents and chemicals.

### Apoptosis assay

UW228-2 cells were seeded at 1 × 10^5^ / well into 6 well plates (Corning; cat CL S3516) and incubated at 37 °C overnight in DMEM containing 10 % FBS. Triplicate wells were cultured in complete media or switched to serum free DMEM for 24 hours before screening with Annexin V APC/PI using the Apoptosis detection kit (Biolegend; cat 640932) according to the manufacturer’s instructions. 30,000 events were acquired using the LSR II flow cytometer with lasers for APC (640 nm laser, emission captured at 660 nm) and PI (488 nm laser with emission captured at 575 nm). Positive controls were generated to inform accurate gating: cells were treated with 200 µM (UW228-2) Etoposide (SIGMA; cat E1383) for 24 hours (Apoptotic cells: Annexin V), or heated at 56°C for 10 minutes prior to labelling (Dead cells: PI).

### Immunofluorescence microscopy

Cells and EVs were immobilised onto CellCarrier 96 well plates (Perkin Elmer; Cat 6005550), fixed with 3.7 % paraformaldehyde, permeabilised with 0.2 % Triton X in PBS and probed using anti-human CD63 antibody (Clone: H5C6, Biolegend Cat: 353005) directly conjugated to FITC and counterstained for polymerised actin using 0.2 × Alexa Fluor 555 Phalloidin (Thermofisher Scientific; cat A34055) and 300 nM DAPI (Biolegend; 422801). Images were captured using the Perkin Elmer Operetta system at x 40 magnification.

### Transmission electron microscopy (TEM)

EVs were immobilised onto ACLAR (poly-chloro-tri-fluoro-ethylene (PCTFE) film) coated with CellTak (Corning: cat 354240) and fixed with glutaraldehyde in sodium cacodylate buffer (pH 7.2) followed by post-fix staining with osmium tetroxide and uranyl acetate (supplied in-house by the FBMH Core Facility). Preparations were dehydrated and embedded in resin to allow serial 60-200 µm sections to be taken. Images were captured using a Biotwin Philips TECNAI G2 transmission electron microscope.

### Cell culture

For each experiment, cells were seeded at 2.5 × 10^6^ cells in 50ml DMEM 10% FBS (complete media) per 225 cm^3^ tissue culture flask and allowed to adhere overnight. On day 2 media was switched to 50 ml serum free DMEM for 24 hours prior to EV isolation and cell preparation. Conditioned media (containing EVs) was removed and the cells washed x 2 with PBS before trypsinisation using 1 × Trypsin-EDTA (Lonza, UK cat: T3924). Cell counts and viability were checked at the time of EV harvest using the trypan blue exclusion assay (0.4% Trypan blue solution; SIGMA; cat T8154).

### Vesicle isolation

Large EVs were harvested using a standard operating procedure (SOP) as previously reported (Figure 1A) (11). For each experiment, EVs were isolated from the serum free, conditioned media from 2 × 225 cm^2^ flasks (pooled; total 100 ml). Cell culture supernatant (conditioned media: CM) was centrifuged in 2 tubes to remove cells (300 × g 5 minutes, × 2) and filtered using a double layered 5µm pore nylon Sieve (Fisher Scientific: BioDesign cat 12994257). The supernatant was collected and centrifuged at 2000 × g for 30 minutes and prepared for ISX analysis. Centrifugation steps were performed using an Eppendorf 5702 bench top centrifuge with an A-4-38 rotor. All EV preparations were performed on the day of analysis and not stored. Experimental procedures for EV isolation have been submitted to the EV-TRACK database (EV TRACK ID: EV190013) (12).

**FIGURE 1:**
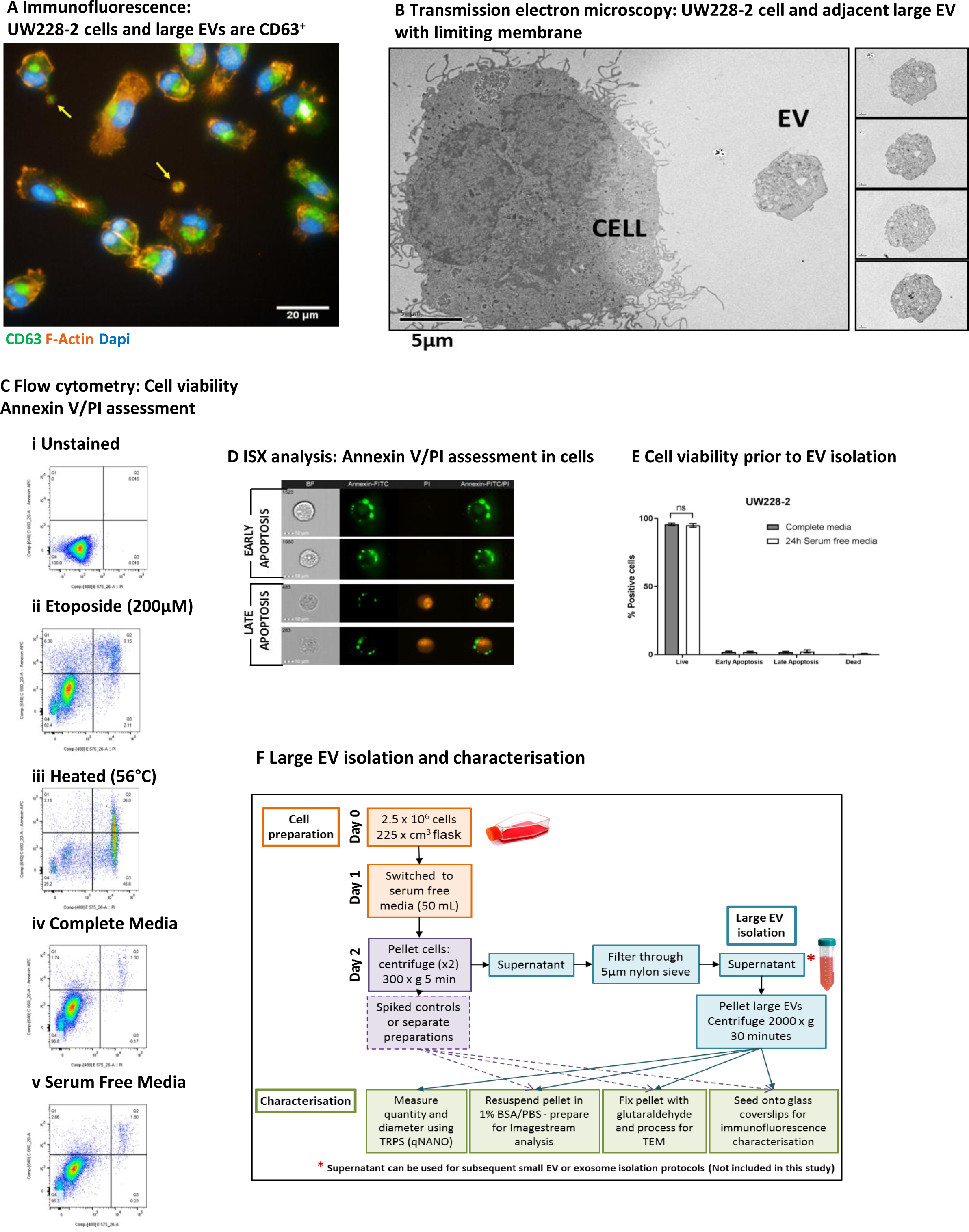
Medulloblastoma cell lines produce large extracellular vesicles (EVs) which can be harvested from serum free conditioned media after 24 hours. **A. Medulloblastoma cells produce EVs *in vitro* which contain polymerised actin and CD63, but no nucleus.** The medulloblastoma cell line UW228-2 was cultured on glass bottom plates for 24 hours; fixed and probed for the tetraspanin and EV marker CD63 (FITC – green). Polymerised actin was labelled using Alexa Fluor 555 phalloidin (f-ACTIN – red). Nuclei were counterstained with DAPI (DAPI – blue). Images of large EVs (yellow arrows) and cells were captured using the Perkin Elmer Operetta system at x40 magnification. The individual images were captured in black and white using the appropriate emission filters and a coloured composite image created using Columbus software. Scale bar represents 20µm. **B. Large EVs have a limiting membrane, contain organelles but no nucleus and are <6µm in size.** Cells and EVs were applied to Poly-d-lysine coated ACLAR film and fixed with glutaraldehyde before processing for Transmission Electron Microscopy. Serial sections were taken to assess sample purity, EV membrane integrity, size and content comparative to the parent cell. Images were captured using a Biotwin Philips TECNAI G2 microscope at x1900 magnification. Scale bar represents 5µm. Left panel shows a cell and a large EV. Right panel: serial sections taken through the entire EV (4 images shown). The wide-field image shows an EV of 5.5µm diameter at its central plane of focus, alongside a parent medulloblastoma cell of 18µm. **C. Cell apoptosis was assessed by flow cytometry**. Unstained cells (i) and cells treated with 200µM Etoposide (ii; Annexin V positive gate) or heated (iii; PI positive gate) provided positive controls which were used to assign accurate quadrant gates to bivariate scatter plots showing PI versus Annexin V APC. Live cells appear in Q4, Annexin only in Q2 (Early apoptosis), dual labelled in Q3 (Late apoptosis) and PI only (Dead cells). UW228-2 cells were cultured for 24h in complete media (iv; DMEM with 10% FBS) or serum free media (v; DMEM only). Representative plots of triplicate experiments shown. **D. Imaging flow cytometry allows visual distinction between early and late stages of apoptosis.** Cells in standard culture conditions were labelled using an Annexin FITC/PI kit and examined using the Imagestream (ISX) Mark II. Images were captured at x60 magnification and representative gallery images demonstrate Annexin V FITC only membrane labelling, indicating early apoptosis (upper panels) and dual Annexin V FITC/PI labelling, indicating late apoptosis (lower panels). **E. UW228-2 cells can be cultured in serum free conditions for 24 hours with no loss in viability.** No difference in viability could be seen between culture conditions: Percentage unlabelled cells (live), Annexin V positive only (Early apoptosis), Annexin V and PI positive (Late Apoptosis) or PI positive only (DEAD) after 24 hours in complete (grey bars) or serum-free medium (open bars). **F. EV isolation protocol from cultured cells was optimised to harvest large EVs with minimal processing.** Cells were seeded into large flasks (225 cm^3^) and allowed to adhere. Media was changed after 24 hours for 50ml serum-free DMEM and cells cultured at 37 °C for a further 24 hours. Trypsinised cells were collected by 2 successive centrifugation steps at 300 × g 5 minutes keeping the supernatant each time. EV containing supernatant was filtered using a double layered 5µm pore nylon membrane by gravity. Filtered, cell-free supernatant was centrifuged at 2000 × g for 30 minutes using a bench top centrifuge with swing out buckets. The resulting cell and EV pellets were processed appropriately for the downstream technique: For flow cytometry cells were processed and analysed separately whilst cells were spiked into wells for direct comparison by microscopy.

### Tunable Resistance Pulse Sensing (TRPS)

Size and quantity was determined by Tunable Resistance Pulse Sensing (TRPS) using the qNano GOLD instrument (iZON Science) as per manufacturer’s instructions. The principles are discussed elsewhere (13). We analysed an aliquot of isolated EVs alongside downstream analyses using overlapping sizes of Nanopores (NP600, NP1000 and NP2000) to provide a full picture of EV size distribution and quantity.

### Cell mask orange labelling

100 mL (2 × 50 ml) EV containing media was used to harvest large EVs for each experiment as described and the 2000 × g pellets re-suspended in either 1) 2ml serum free DMEM or 2) 2 mL Cell Mask Orange (CMO: 2.5 µg/mL in serum free DMEM) (Thermofisher Scientific cat C10045) and both were incubated at 37° C for 10 minutes.

### Antibody labelling

See table 1 for antibody details and manufacturer information. Antibody titrations were performed for each antibody using parent cells and EVs. In all cases, the maximum recommended volume (5 µl) provided the greatest fluorescence signal from the EVs. Antibody only controls (no EVs) were included in the ISX analysis and showed no fluorescence events above the unstained gate in each case. Both CMO labelled and unlabelled EVs preparations were washed by addition of 5 mL 1% BSA/PBS and centrifuged at 2000 × g for 30 minutes. The pellets were re-suspended in 700 µl 0.2 µm filtered 1% BSA/PBS and split into 7 × 100 µl aliquots. For each labelling combination, 5 µl directly conjugated primary antibodies were added simultaneously as follows: Non-CMO labelled EVs were used for unstained, single labelled CD63 BV421, CD9 AF647 or LAMP1 AF647 and CMO FMO controls (x2) (BV421 + AF647: both antibodies were screened). The final aliquot was used to establish EV concentration and diameter range using TRPS analysis on the qNANO. The CMO labelled EV pellet was re-suspended in 700 µl 0.2 µm filtered 1% BSA/PBS and split into 7 × 100 µl aliquots. CMO labelled EVs were used as single stained (CMO+ only), AF647 FMO (CMO + BV421), BV421 FMO (CMO + AF647), and multiplexed CMO + CD63 BV421 + either CD9 AF647 or LAMP1 AF647. Antibodies were incubated for 1 hour on ice in the dark. EVs were washed by addition of 500 µl 0.2 µm filtered 1% BSA/PBS and centrifuged at 2000 × g for 30 minutes. Resulting pellets were re-suspended in 75 µl 0.2 µm filtered 1% BSA/PBS for ISX analysis. Fully labelled EV preparations were treated post-acquisition with 0.1% Triton-X 100 and acquired again to demonstrate EV lysis and loss of signal.

### Imagestream Acquisition

Sheath buffer (PBS without calcium and magnesium: SIGMA D5652) was filtered using 0.2 µm bottle top filters (Nalgene: FIL8184) to minimise background signal. Internal instrument calibrations were performed before every run according to manufacturer’s guidelines using the ASSIST Calibrations to include: camera synchronisation, spatial offsets, dark current, bright field crosstalk coefficient, core stage position, horizontal laser, side scatter and a retro illumination scheme to maximise the amount of light incident. This was followed by a series of internal operations designed to measure performance including excitation laser power, bright field uniformity and focus. Specific laser powers used for this study are detailed in Table 2.

**TABLE 2:**
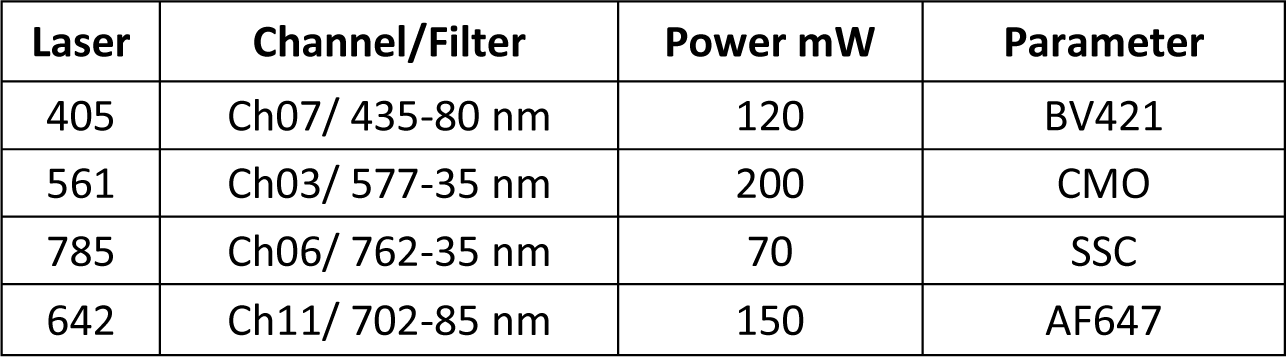
Settings used for Imagestream.

Speed beads with an exaggerated irregular surface were incorporated into every analysis for internal calibration. A dedicated laser was assigned to assess side scatter (CH 06; SSC: 785 nm laser). For each experiment, a separate readout was obtained from 0.2 µm filtered 1% BSA/PBS acquisition buffer alone. All events were acquired for 5 minutes and visualised using bivariate plots of side scatter against fluorescence intensity. Details of the gating hierarchy which formed the final EV analysis template are shown in Figure 4.

For cells; 10,000 – 30,000 total events were acquired. EVs were acquired for 5 minutes at x 60 magnification using lasers as described (Table 2) (Image stream, Amnis, Seattle, United States). The 60 x objective provides a Numerical Aperture of 0.9 enabling resolution of 0.3 µm^2^/ pixel (14).

### COMPENSATION

Antibody labelled compensation beads (anti-mouse compensation beads: BD Biosciences cat 552843) were used to acquire single colour controls within the channels used for this study. The final compensation matrices was constructed by the wizard (INSPIRE) with manual adjustment in consultation with the manufacturer’s specialist adviser and applied to the .rif files of all controls, dual and triple labelled EVs. Data were analysed using the IDEAS software (version 6.2). The compensation matrices and analysis template were applied using the batch processing tool to all .rif files to produce .daf files for each sample. FCS files were exported and uploaded onto the Flow Repository according to the requirements.

### MASK SELECTION FOR ASSESSMENT OF EV DIAMETER

We investigated which of the diameter masks available within the IDEAS software would be most accurate for EVs. We used non-fluorescence size calibration beads (Thermofisher: F13838) to validate the masks. The beads were acquired using the same laser powers and settings as the EV preparations and analysis templates were constructed to identify which mask fitted most closely to the bead diameter according to the manufacturer’s instructions. We found that applying the diameter mask Erode (03; indicating 3 pixel erosion) to the bright field channel most closely assigned the correct diameter. This mask formed part of an analysis template which was applied to all samples using the batch analysis tool.

### MESF CALCULATION

To enable comparisons between experiments, Molecules of Equivalent Soluble Fluorochrome (MESF) values were calculated as previously described (14, 15). Quantibrite PE beads (Cat: 340495. Lot: 90926) were the closest available calibration beads for the fluorescent channel used to detect Cell Mask Orange labelled EVs. A fresh aliquot of lyophilised Quantibrite beads was reconstituted for each run, and 5000 events were acquired using the identical laser settings for each fluorophore as described. The SSC laser (channel 06) was adjusted to ensure the beads could be visualised on the bivariate plots and therefore each bead could be gated as a separate population and the median fluorescence intensity recorded. The CMO ^+^ events were then analysed for expression of the EV markers included in this study: CD63, CD9 and LAMP1.

## RESULTS

**Large EVs from the medulloblastoma cell line UW228-2 are visible with light microscopy, can be isolated from viable, serum-free cell culture supernatant with intact membranes, and contain a polymerised actin cytoskeleton. A proportion of these large EVs express reported EV markers.**

In this study the large EV isolation SOP was applied to the SHH-driven medulloblastoma cell line UW228-2. In accordance with the recently published international consensus MISEV2018 guidelines (1), we demonstrated the existence and membrane integrity of large EVs using both light and electron microscopy (Fig 1A and B). The pan-EV marker CD63 was expressed by parent medulloblastoma cells and a proportion of large EVs in viable cultures (Figure 1A). Large EVs also exhibited an active cytoskeleton indicated by the polymerised actin marker Phalloidin, but did not stain for DAPI, indicating that they did not contain nuclear double-stranded DNA. EV membrane integrity, size and intra-vesicle content were examined by transmission electron microscopy (TEM) (Figure 1B). Isolated EVs were spiked with medulloblastoma parent cells for comparison and adhered to ACLAR film coated with CellTak prior to TEM. Serial sections demonstrated that the EVs had a limiting membrane, internal organelles but no nucleus, and were independent from cells. By contrast, cells had internal organelles including a nucleus and cytoplasmic protrusions indicative of filopodia.

Cells were grown in serum-free medium for 24 hours prior to EV isolation, staining and analysis. Experimental conditions were optimised to eliminate false positive membrane labelling or bovine EV contamination (16). To assess whether growth in serum-free medium resulted in increased cell apoptosis, phosphatidyl serine exposure on the surface of cells was examined. Figure 1C indicates the quadrant gating strategy applied using positive controls and the comparison of cells grown in standard culture versus serum free conditions. Imaging flow cytometry was also performed to visualise Annexin V/PI staining on parent cells (Figure 1D). In triplicate experiments, there was no significant difference between the viability of cells cultured in complete or serum free media (Figure 1E). Figure 1F provides a workflow for the isolation and characterisation of large EVs used in this study. At the time of each EV isolation, total parent cell count and viability was assessed using trypan blue exclusion and found to be 7.9 × 10^6^ cells (98% viable), 8.3 × 10^6^ cells (99% viable) and 6.8 × 10^6^ cells (98% viable).

### EVs are highly heterogeneous and differentially express EV markers

A reliable fluorescence marker was essential to demarcate EVs from background scatter events. Cell mask orange (CMO) is a fluorescent plasma membrane label composed of amphipathic molecules comprising a lipophilic moiety for membrane loading and a negatively charged hydrophilic dye for anchoring of the probe in the plasma membrane. We performed a titration using the parent cells with serial dilutions of the dye in serum free media ranging from 1 in 1000 (5 µg/ml) to 1 in 100 000 (50 ng/ml) (Figure 2 A) and found that 2.5 µg/ml (1 in 2000) was an optimal concentration providing high median fluorescence intensity without saturation. Using the Imagestream Mark II (ISX) we confirmed that the final concentration of CMO labelled EVs in a typical preparation did not result in coincidence events or swarm which could lead to false positive results when looking at multi-colour labelling. Serial dilutions of CMO labelled EVs showed a linear decrease in objects per ml (Figure 2 B i) with an increasing dilution factor, whilst the fluorescence intensity remained stable across the dilutions (Figure 2 B ii). To facilitate reproducible reporting across experiments and to offer a means to standardise EV measurements, we determined the minimum MESF value for CMO+ events which would distinguish between unstained and CMO labelled EVs using our staining protocol, as described elsewhere (14). Figure 2 C shows a representative bivariate plot (i) of the low, medium-low, medium-high and high fluorescence bead populations. The histogram (ii) was used to determine the MFI of each peak and log fluorescence intensities were converted to MESF values using information supplied by the manufacturer (iii). Linear regression of log MFI versus log MESF (iv) was used to calculate the MESF corresponding to the minimum fluorescence intensity of events within the designated CMO ^+^ gate as follows. The maximum MESF values of the unstained EVs in each experiment were : Replicate 1: 271.42 = MESF 48.23; Replicate 2: 271.0 = MESF 49.27; Replicate 3: 272.88 = MESF 42.48. Therefore, to set a standardised lower threshold of detection which could be used across replicate experiments we assigned a lower MESF threshold for CMO+ events of 50 (Example shown in Figure 2 C v). We therefore report here CMO ^+^ events as number of events > MESF 50.

**FIGURE 2:**
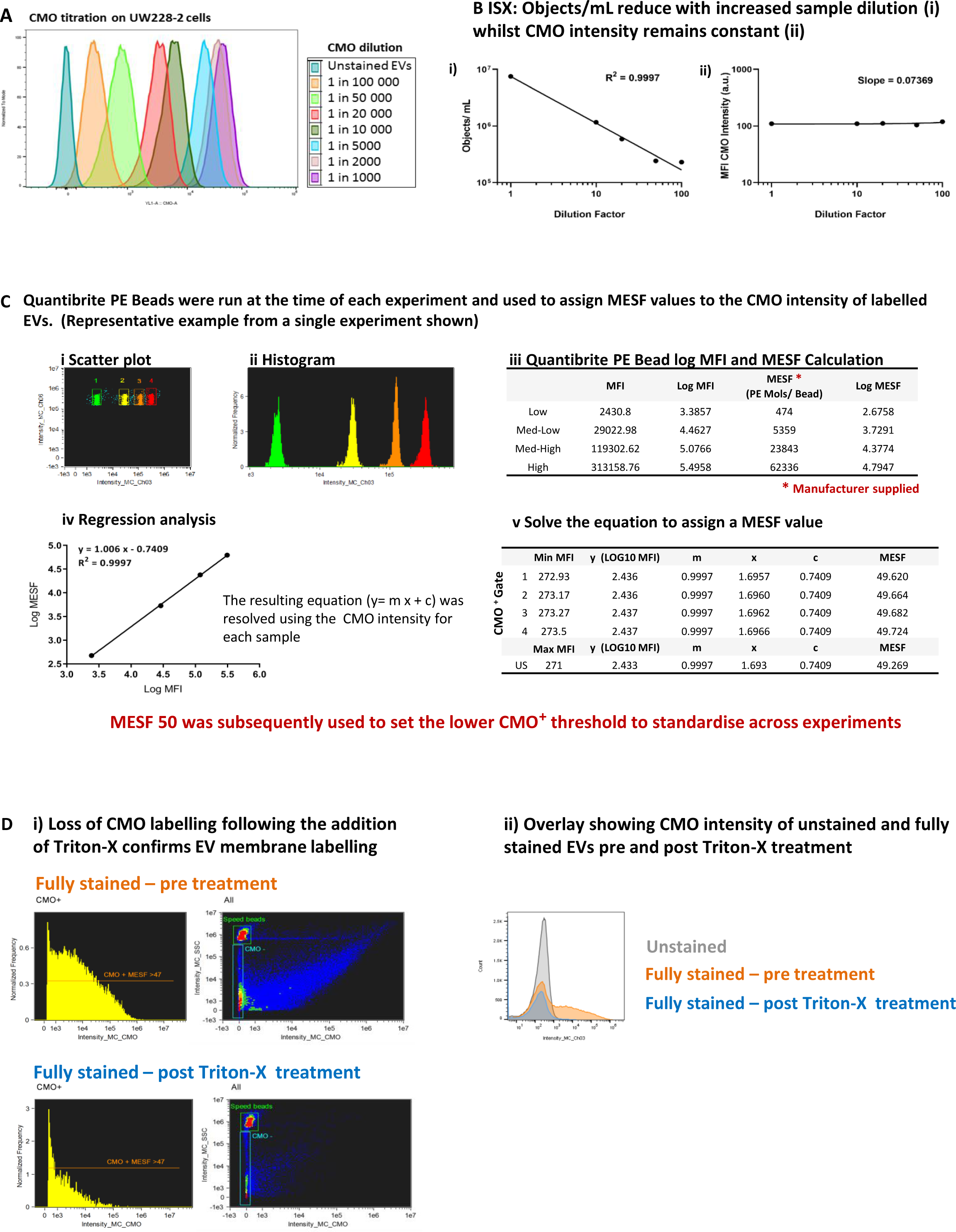
Cell Mask Orange can label EVs to enable standardisation across experiments. **A. Cell Mask Orange (CMO) labelling was optimised by titration using parent cells.** Serial dilutions from 1 in 1000 – 100 000 were used to label parent UW228-2 cells and analysed by flow cytometry. A clear relationship between CMO concentration and fluorescence intensity is shown. **B. Sample dilution was used to validate EV staining protocol.** Using a CMO dilution of 1 in 2000 (2.5 µg/mL), the Imagestream acquired fewer CMO positive objects per ml with increasing dilution of CMO labelled EVs (i), whilst the CMO fluorescence intensity in channel 03 was maintained (ii). **C. Quantibrite PE beads were acquired at the time of each experiment to assign MESF value.** Quantibrite PE beads were separated on a bivariate plot of intensity against side scatter (i). The fluorescence intensity of each bead set was gated on the histogram (ii) and assigned a MESF value according to the number of PE molecules per bead as provided by the manufacturer (iii). Log MFI against Log MESF provided a standard curve for each acquisition (iv). Regression analysis of was used to extrapolate the equivalent MESF value for CMO intensity for unstained and labelled samples (v) which enabled lower threshold for CMO intensity to be set to distinguish between unstained and CMO labelled EVs. Representative experiment shown. **D. CMO labelling was diminished following the addition of detergent.** CMO positive (CMO **^+^**) EVs could be distinguished from unstained EVs by ISX (i). Treatment of the same sample with Triton-X 100 diminished the CMO labelling (ii). Exported .fcs files from the same acquisition were overlaid using FlowJo 10.6. The fully stained sample (orange) shows increased fluorescence intensity in channel 3. The same sample post Triton–X treatment (blue) showed a reduction in fluorescence intensity to a similar level of the unstained sample (grey).

Previous reports have identified that lipid dye aggregates can mimic EVs when using fluorescence lipid markers (17). We used Triton-X 100 treatment (18) of fully labeled EVs to disrupt EV signals demonstrated by a loss of CMO + events (Figure 2 D i). Figure 2 D (ii) shows an overlay of unstained EVs, fully stained EVs and fully stained EVs treated with Triton-X 100. The MFI of the post-Triton-X 100 treated sample is reduced to the level of the US-EVs.

### Large EVs size distribution and quantity was assessed using Tunable Resistance Pulse Sensing (TRPS)

At the time of each preparation, an aliquot of freshly isolated EVs was assessed using the qNano GOLD particle counter to ascertain EV size distribution (diameter) in terms of percentage population (%) and concentration (particles/ mL) prior to labelling for ISX analysis. Using a series of 3 overlapping Nanopores (Figure 3 A i) each with an optimal size range which span a total size of approximately 275 nm to 5.7 µm, we assessed the size range of particles in the EV preparations. Representative profiles of diameter against percentage population are shown for a single sample using NP600, NP1000 and NP2000 Nanopores (Figure 3 A ii-iv). By selecting overlapping Nanopore sizes, the same calibrator beads (1000 nm) could be used and therefore it was possible to overlay the resulting profiles to visualise the total large EV population within each sample (Figure 3 A v). As expected, the EVs present in each biological replicate were heterogeneous in size. However, the size range of EVs across biological replicates was consistent (250 nm to 6 µm); with some variation in the median diameter per sample. Size is presented with a bin of 100 (nm) and in each case; the most prevalent large EVs were detected using the NP600. The median size across the replicates using the NP600 for comparison were 250-350 nm 5.15 × 10^7^; 450-550 nm 1.7 × 10^7^ and 250-350 nm 2.8 × 10^8^. The starting volume of EV-containing media used per replicate was 100 ml. The resulting pellet was re-suspended in a total of 700 µl PBS (prior to labelling). As an example, applying this dilution factor (142.86) to the first replicate equates to 7.4 × 10^9^ EVs specifically of the 250-350 nm size range that were released by 7.9 × 10^6^ UW228-2 cells (NB cell count at harvest) in 24 hours into 100 ml serum free media. It should be noted that due to the number of washes and centrifugation steps during the isolation, this can only be considered as an illustration of the quantity of EVs produced. The total larger EVs as counted using the NP2000 (935 nm – 5.7 µm) were less common but are nevertheless abundantly present in concentrations of 2.3 × 10^8^, 6.9 × 10^8^ and 2.1 × 10^9^ in the 100ml harvested media from this particular cell line. The total particle count for each Nanopore size, across 3 biological replicates is shown in Table 3. It should be noted that the counts shown are not cumulative as they represent the same sample analysed across 3 Nanopore sizes.

**FIGURE 3:**
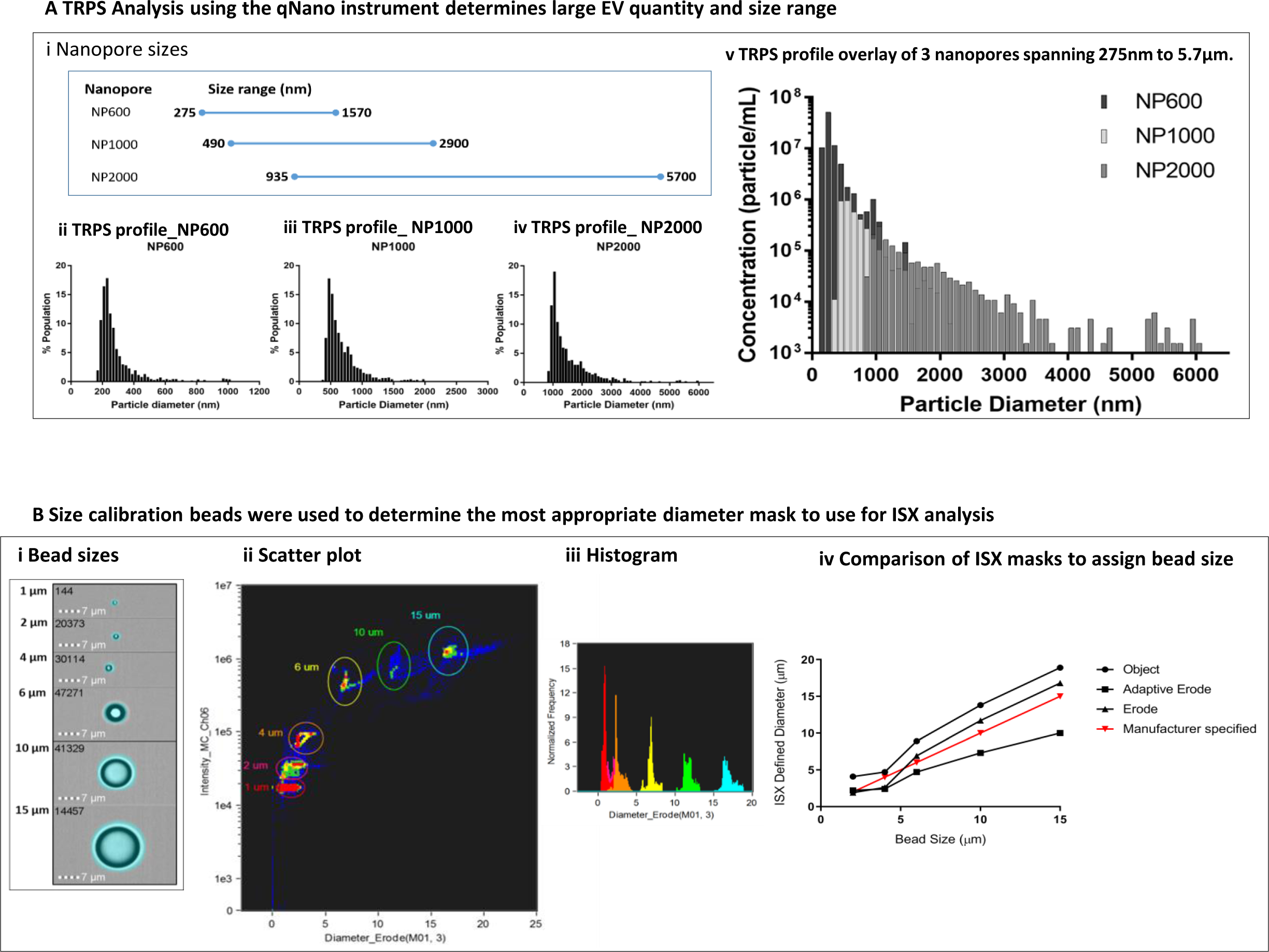
EV size and quantity was assessed by Tunable Resistance Pulse Sensing (TRPS) using the qNANO instrument (iZON Science) and compared to diameter masks in the ISX IDEAS software. A. Three Nanopores of over-lapping size ranges were used to determine the particle diameter and concentration in each EV preparation (i). Size profiles were established for each individual Nanopore (ii NP600; iii NP1000; iv NP2000). The overlaid profiles of diameter by concentration provided by the 3 Nanopores show a size range of 250nm and 6µm against a common calibrator bead of 1000nm (representative sample shown) (v). **B. Calibration beads of known size were used to determine the most accurate diameter mask.** The masks available within the ISX IDEAS software for diameter analysis were compared. The 3 which assigned the most accurate sizes according to manufacturer specified beads sizes were the Object, Erode and Adaptive Erode masks. These were compared across bead size by applying the mask (ii) visualising a scatter plot (iii) and histogram for each mask type. For the size range within our EV preparations, we found the Erode mask to be the most accurate. (iv)

**FIGURE 4:**
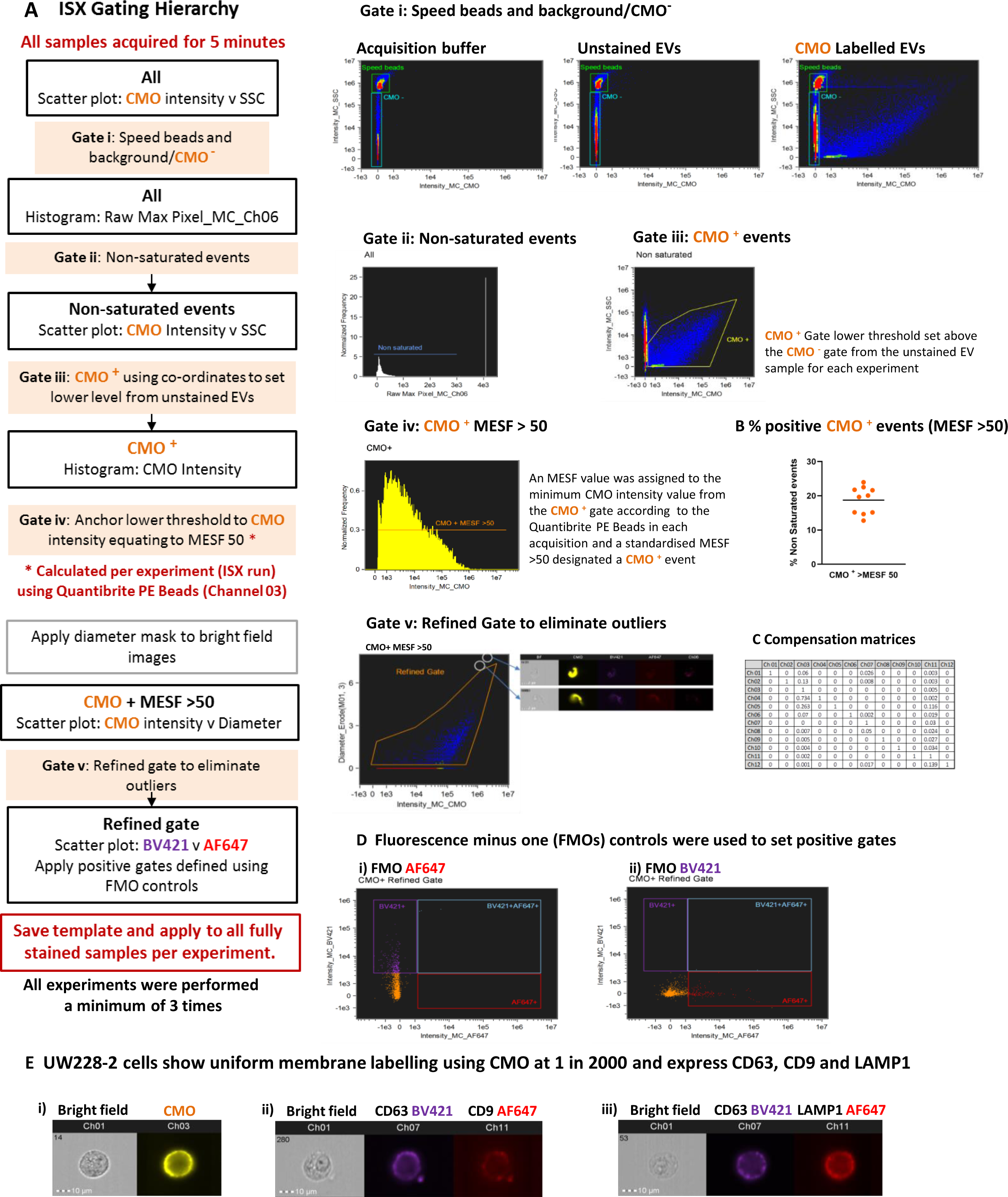
Optimised acquisition by Imaging flow cytometry and florescence membrane labelling can distinguish large extracellular vesicles from background and enable multiplex labelling. **A. Hierarchical gating can confidently refine the EV population for characterisation.** Samples were acquired for 5 minutes using the ISX INSPIRE software and all events visualised using bivariate plots for fluorescence intensity in channel 03 (CMO) and channel 06 (side scatter). Initial gates for speed beads and CMO ^−^ events were set using the Acquisition buffer only and unstained EV samples included in each run (Gate i). These gates were applied to the labelled samples. Saturated events were excluded from the analysis by gating on the histogram plot: Raw Max Pixel for Channel 06 (Gate ii) thus removing the speed beads and very high side scatter events. An initial CMO+ gate was placed (Gate iii). A lower fluorescence threshold of MESF >50 in channel 3 (CMO) was set in each experiment using the regression analysis of Quantibrite PE Beads as described. The CMO + gate was anchored using co-ordinates for the equivalent fluorescence intensity and a further CMO + MESF >50 gate applied to the histogram (Gate iv). Finally a refined gate was used to eliminate outliers (Gate v). The analysis template was set for each experiment and applied to all samples. **B. Percentage positive CMO^+^ events (MESF >50).** The percentage of CMO+ events (MESF >50) is represented by an orange dot for 10 replicate experiments. Black bar represents mean 18.73% (+/− 1.2 SEM). **C. A compensation matrix was applied to all samples.** The compensation wizard was used to create a matrix which was applied to all samples. **D. Single, double and triple labelling protocols were used for compensation and accurate gating.** Dual labelled EVs provided Fluorescence minus one (FMO) controls for **(i)** AF647; EVs labelled with CMO and CD63 BV421) or **(ii)** BV421; (EVs labelled with CMO and LAMP1 AF647) and used to set positive gates. **E. UW228-2 cells were used to optimise multiplex labelling.** Cell mask orange (CMO) provides a general membrane label (i) and could be used to co-label with EV markers CD63 BV421 and CD9 AF647 (ii) or CD63 BV421 and LAMP1 AF647 (iii). Representative gallery images from IDEAS software are shown.

**TABLE 3:** Total particle count using different Nanopore sizes (qNANO) in biological replicates.

|  | NP600 | NP1000 | NP2000 |
| --- | --- | --- | --- |
| **1** | $8.4 \times 10^8$ | $3.8 \times 10^6$ | $1.6 \times 10^6$ |
| **2** | $4.9 \times 10^7$ | $1.4 \times 10^7$ | $4.8 \times 10^6$ |
| **3** | $5.0 \times 10^8$ | $4.4 \times 10^7$ | $1.5 \times 10^7$ |

Size calibration beads were used to determine the most appropriate diameter mask to use for ISX analysis of EV diameter. The bead sizes provided by the manufacturer were 1, 2, 4, 6, 10 and 15 µm. We used the feature tool in IDEAS to apply a range of different diameter masks to the bright field image of the beads (Figure 3 B i). We selected bright field for this using non-labelled beads because the Quantibrite beads showed over exaggeration of EV size, possibly due to saturation and flare of fluorescence. The beads were visualised on bivariate plots of diameter versus side scatter (channel 06). Density plots (Figure 3 B ii) allowed the bead populations to be gated individually and subsequent histograms (Figure 3 B iii) to be viewed. We compared the diameters for each bead population assigned using the Object, Adaptive Erode and Erode Masks against the manufacturer specified size (Figure 3 B iv) and found the Erode mask on the bright field image, using 3 pixel reduction to be the most comparable.

### Fluorescence membrane labelling can be used to distinguish EVs from background and speed beads

We devised an analysis template within IDEAS which comprised a hierarchical gating strategy aimed at characterising heterogeneous large EVs (Figure 4 A). All buffers were filtered using a 0.2 µm filter and samples were acquired for 5 minutes to avoid recording different amounts of background per acquisition. Speed beads were included. The speed beads and CMO ^−^ events were defined using density plots of channel 03 (CMO) versus channel 06 (side scatter) intensity (Gate i). Running acquisition buffer only (left side) and unstained EVs (centre) showed the instrument detected a high level of background with low to medium side scatter. Labelling with the cell membrane dye (CMO) therefore helped to distinguish between the low CMO intensity/ low side scatter EVs and background detected in both the acquisition buffer and unstained sample (right side). Applying a gate to capture low Raw Max Pixel events (channel 06 – SSC: Gate ii) eliminated those events with saturated side scatter, including the speed beads. Outliers in the side scatter versus CMO intensity plots were individually inspected and found to be dual events comprising a speed bead (CMO-, high SSC) and an EV (CMO+, low SSC) which occupied the flow chamber at the same time. This resulted in aberrant events with both high CMO intensity and high SSC and was therefore excluded from further analysis.

Events classed as CMO^+^ at this point were included in the initial CMO^+^ gate (Gate iii). Within the same acquisition, we ran the Quantibrite PE beads which enabled a minimum threshold for CMO^+^ events to be calculated and converted to MESF units, as described. The lower MESF cut off for this experiment was defined at > 50 and a further gate on the CMO intensity histogram (Gate iv) captured all CMO^+^ events with an MESF >50 for downstream analysis. The proportion of CMO^+^ events as a percentage of all events acquired over the 5 minute period varied across 10 replicates (mean 18.73% +/− 1.2 SEM) CMO^+^ MESF > 50; Figure 4 B). A final gate was included to eliminate outliers which were either high CMO intensity but appeared on the bright field image to be membrane fragments (Gate v) or were not assigned a diameter due to lack of bright field image focus. This stringent gating strategy was designed to eliminate any events which could not be further analysed for EV marker expression or diameter.

We tested a number of approaches to apply compensation to our data in consultation with the manufacturer’s specialists. Initially, we attempted to construct a matrix using single stained EVs; however, the single pixel fluorescence was insufficient for the in-built compensation Wizard to assign a matrix. Next, we tried single stained parent cells, but the compensation matrix formulated by the wizard resulted in over-compensation and negative fluorescence intensity events. Over-compensation was thought to be due to the imbalance between the strong fluorescence signal resulting from cell mask orange, a membrane marker which indiscriminately labels lipids, and relatively weak fluorescence signal from target specific antibodies. These experiments were originally performed using a FITC conjugated CD63 antibody, however we found the level of adjustment required between channel 02 (FITC) and channel 03 (CMO) contributed to the negative populations. We re-optimised using the same CD63 antibody clone (HSC6) conjugated to BV421 (channel 07) which is spectrally more distinct from CMO. Finally, we found that using commercially available compensation beads labelled with our antibodies and acquired in the channels used for our study, in conjunction with assisted manual adjustment, provided the most reproducible compensation matrix.

We combined CMO with BV421 and AF647 and Fluorescence Minus One (FMO) controls for each fluorophore were used to set positive gates (Figure 4 D). The parent UW228-2 cells were screened for expression of the EV markers chosen for this study: CD63, CD9 and LAMP1 as defined in the latest MISEV guidelines (1). Multiplex labelling was performed using the following combinations: CMO with CD63 BV421 and CD9 AF647; or CMO with CD63 BV421 and LAMP1 AF647.

### Multiplex labelling reveals heterogeneity in EV marker expression

The proportion of CMO^+^ only events varied across the 10 replicates (Figure 5 A). The mean percentage positive CMO^+^ (MESF >50) events (63.3% +/− 4.6 SEM) which did not demonstrate expression of any EV markers included in this study were designated CMO only. Considering the single EV markers, the proportion of EVs positive for each EV marker varied CD63 19.0%, (+/− 2.2 SEM), CD9 mean 10.3% (+/− 0.5 SEM); LAMP1 47.6% (+/− 3.8 SEM). These data shown LAMP1 to be the most prevalent EV marker screened in this study.

**FIGURE 5:**
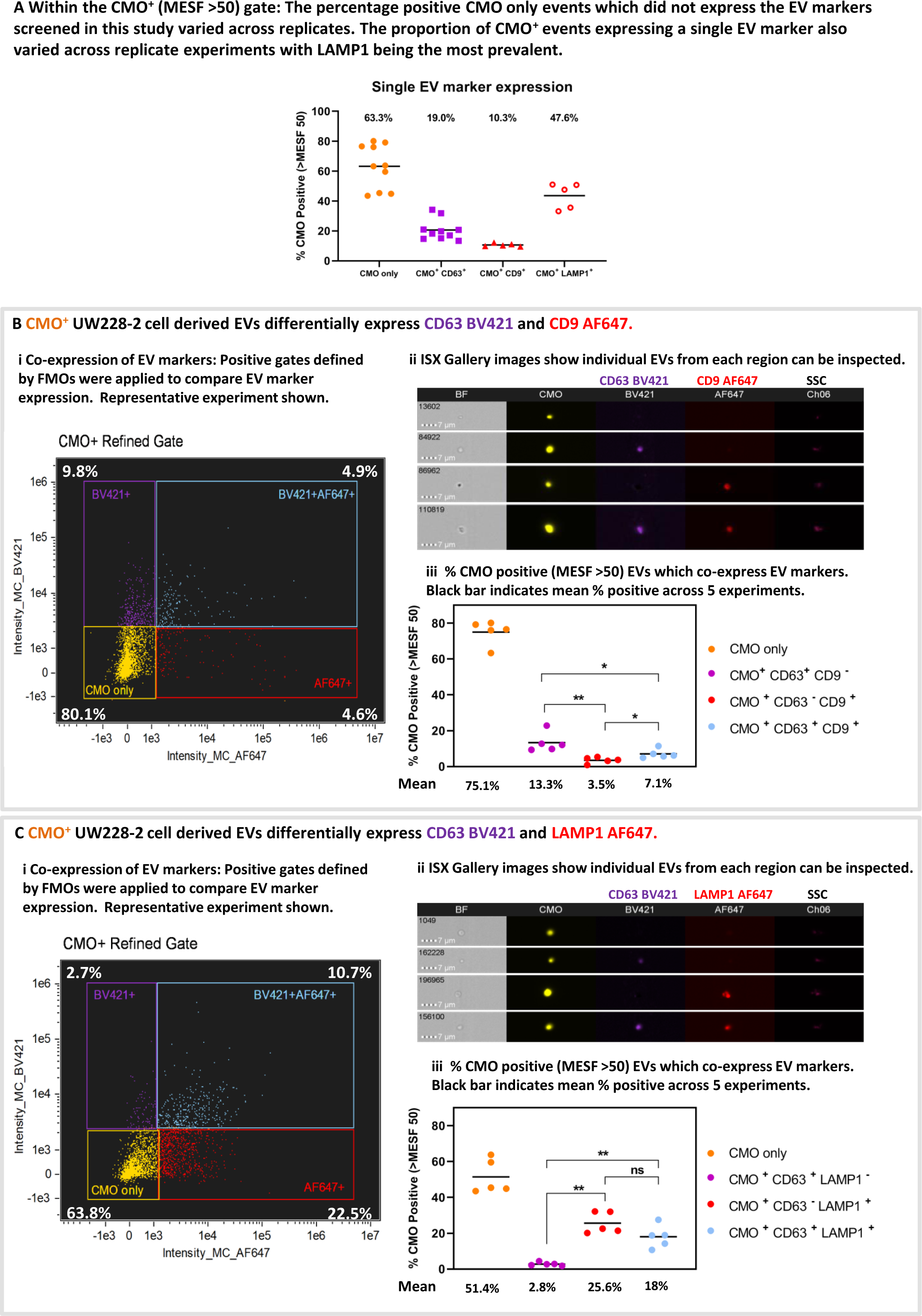
Isolated large EVs can be triple-labelled and individual events can be scrutinised post-acquisition. **A. The proportion of CMO^+^ events expressing a single EV marker varied across replicates.** The percentage positive events within the CMO^+^ MESF >50 gate which expressed either no EV marker (CMO only), CD63, CD9 or LAMP1 varied across replicates. Black bar represents the mean across 10 (CMO or CD63) or 5 (CD9 or LAMP1) replicates. CMO only 63.3% (+/− 4.6 SEM), CD63 only 19% (+/− 2.2 SEM), CD9 (+/− 0.5 SEM) only 10.3% and LAMP1 only 47.6% (+/− 3.8 SEM). **B. UW228-2 cell derived EVs differentially express EV markers CD63 and CD9, or both**. EVs were labelled with CMO, CD63 BV421 and CD9 AF647. CMO ^+^ events (MESF >50) were analysed for co-expression with CD63, CD9 or both. Quadrant gating of bivariate plots for fluorescence intensity in channel 07 (BV421) and channel 11 (AF647) demonstrated single labelled (CMO ^+^ only), dual and triple labelled EV populations (i). Percentage positive in each quadrant is shown. Representative experiment. Gallery images display examples of individual events from each of the 4 quadrants (ii). Bright Field (BF) and side scatter (SSC) channels are shown alongside CMO, CD63 BV421 and CD9 AF647. Of the CMO^+^ MESF >50: 75.1% (+/− 3.0 SEM) were CMO ^+^ only; 13.3% (+/− 2.4 SEM) expressed CD63 BV421 and 3.5% (+/− 0.8 SEM) expressed CD9 AF647. 7.1% (+/− 1.1 SEM) of the CMO ^+^ events expressed both CD63 and CD9. **C. UW228-2 cell derived EVs differentially express EV markers CD63 and LAMP1, or both**. EVs were labelled with CMO, CD63 BV421 and LAMP1 AF647. CMO ^+^ events were analysed for co-expression with CD63, LAMP1 or both. Quadrant gating of bivariate plots for fluorescence intensity in channel 07 (BV421) and channel 11 (AF647) demonstrated single labelled (CMO ^+^ only), dual and triple labelled EV populations (i). Percentage positive in each quadrant is shown. Representative experiment. Gallery images display examples of individual events from each of the 4 quadrants (ii). Bright Field (BF) and side scatter (SSC) channels are shown alongside CMO, CD63 BV421 and LAMP1 AF647. Of the CMO^+^ MESF >50: 54.1% (+/− 4.3 SEM) were CMO ^+^ only; 2.8% (+/− 0.4 SEM) expressed CD63 BV421 and 25.6% (+/− 2.7 SEM) expressed LAMP1 AF647 whilst 18.0% (+/− 2.8 SEM) of the CMO ^+^ events expressed both CD63 and LAMP1. (* *P* = 0.0159 or 0.0317; ** *P* = 0.0079). Mann Whitney U test.

Differential expression and co-expression of EV markers was examined between five replicate experiments. Representative bivariate plots of AF647 intensity against BV421 intensity (Figures 5 B i and C I for CD63/CD9 and CD63/LAMP1 respectively), and representative galleries (Figure 5 B ii and C ii) of EVs displaying each labelling combination taken from the quadrant plots are shown. The proportion of EVs which co-express EV markers within each gate for the five replicates is shown in Figure 5 B iii and C iii). Where the EVs were labelled for CMO, CD63 BV421 and CD9 AF647 (Figure 5 B iii): 75.1% (+/− 3.0 SEM; yellow) were CMO only, 13.3% (+/− 2.4 SEM; purple) were CMO ^+^ CD63 ^+^ CD9 ^−^; 3.5% (+/− 0.8 SEM; red) were CMO ^+^ CD63 ^−^ CD9 ^+^ and 7.1% (+/− 1.1 SEM; blue) were CMO ^+^ CD63 ^+^ CD9 ^+^. There were significantly fewer CD63 ^−^ CD9 ^+^ EVs compared with CD63 ^+^ (*P* = 0.0079) only or CD63 ^+^ CD9 ^+^ EVs (*P* = 0.0159). Similarly there were fewer EV co-expressing CD63 and CD9 compared with CD63^+^ alone (*P* = 0.0317). (Mann Whitney U test). Considering the EVs labelled with CMO, CD63 BV421 and LAMP1 AF647 (Figure 5 C iii). 51.4% (+/− 4.3 SEM; yellow) were CMO^+^ only, a comparatively low 2.8% (+/− 0.4 SEM; purple) were CMO ^+^ CD63 ^+^ LAMP1 ^−^; whilst 25.6% (+/− 2.7 SEM; red) were CMO ^+^ CD63 ^−^ LAMP1 ^+^ and 18% (+/− 2.8 SEM; blue) co-expressed both CD63 and LAMP1. There were significantly more EVs which expressed LAMP1 compared with CD63 alone (*P* = 0.0079) and more expressing both CD63 and LAMP1 compared with CD63 alone (*P* = 0.0079).

### ISX allow correlative analyses of diameter with EV marker expression for phenotyping

We applied the Diameter Mask; Erode (M01, 3) in IDEAS software to all fully labelled replicates using the Batch analysis function and compared the range of diameters present within the CMO^+^ MESF >50 gate. Figure 6 A shows single event diameters within the 10 replicates. This median diameter (indicated with the black line) varied significantly across the replicates (922 – 1129nm: Kruskal-Wallis; *p > 0.0001*). However, these data represent EVs isolated as 3 biological replicates and when each replicate is analysed using ANOVA there was no significant difference between the median diameters of each EV preparation. There was a clear correlation between EV diameter and CMO intensity as shown in Figure 6 B. Regression analysis was performed on the individual exported feature values from each replicate (2 representative plots are shown from different biological replicates). (*p* < 0.0001)

**FIGURE 6:**
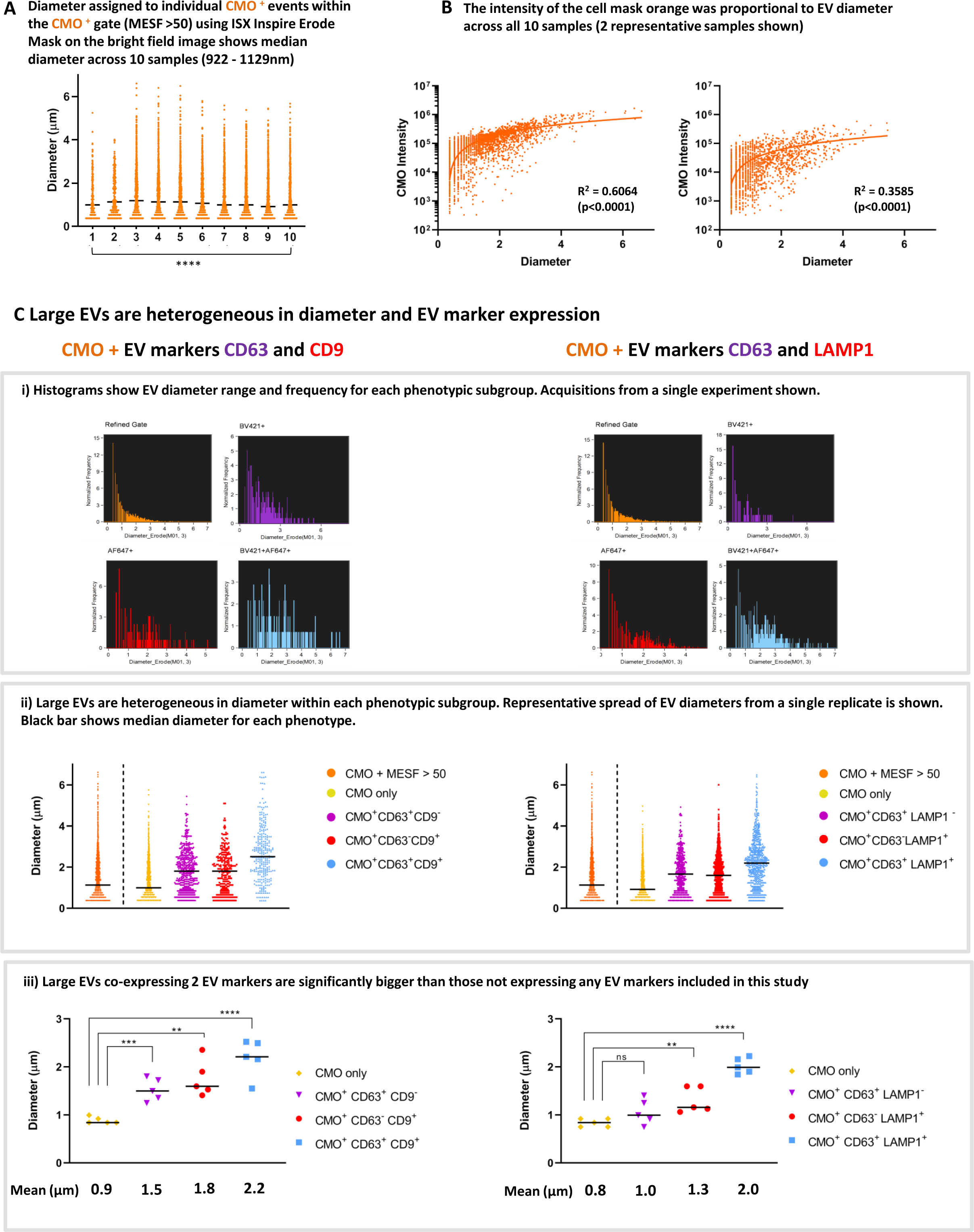
The ISX can accurately assign diameter to large EVs and facilitate individual event analyses to phenotype heterogeneous EV populations. **A. The median diameter of CMO ^+^ EVs from UW228-2 cells was comparable across 10 samples.** The erode mask was applied to the bright field images of those EVs which were included in the CMO ^+^ >MESF 50 gate. The diameter of individual events within this gate was exported from IDEAS into PRISM for analysis. This figure shows a median diameter (represented by the black bar) of 922 – 1129nm across the 10 samples and visualises the broad size range within each sample. (**** *p > 0.0001* Kruskal-Wallis test). **B. The fluorescence intensity of the CMO was proportional to EV diameter.** Bivariate plots showing CMO intensity against diameter consistently demonstrates a proportional relationship between EV size and intensity of the membrane label across samples run in triplicate experiments. 2 representative samples shown. (Regression analysis *p < 0.0001*) **C. Comparison of EV phenotypes across triplicate experiments.** Individual histograms displaying EV diameter for each phenotype demonstrated differential size ranges according to EV marker(s) expression (i a and i b). In each case, those labelled with CMO only and not expressing either CD63, CD9 or LAMP1 have a smaller median diameter compared with EVs expressing CD63 and/or CD9 or LAMP1 (ii a and ii b). In both cases, those EVs co-expressing both CD63 with CD9 (iii a) or CD63 with LAMP1 (iii b) are significantly larger than CMO only. (\*\**p* < 0.005, *** p < 0.0004, **** p < 0.0001)

We found that large EVs are heterogeneous in both diameter and EV marker expression. Histogram plots of individual events exported from the quadrant positive gates show the different diameter range and frequency within each phenotypic subgroup (Figure 6 C i). Representative acquisitions from a single replicate are shown. ISX analysis facilitates the comparison of individual EV diameters from within each phenotype (Figure 6 C ii). A representative plot for all CMO ^+^ (MESF > 50) in a single replicate of each phenotype is shown and the black bars represent the median diameter for each subgroup. Within this typical replicate, the median diameter and range for each subgroup was as follows: All CMO >MESF 50 1.1 µm (362 nm – 6.2 µm), CMO only 1.0 µm (362nm – 5.4 µm), CMO^+^ CD63^+^ CD9^−^ 1.8 µm (362 nm – 5.1 µm), CMO^+^ CD63^−^ CD9^+^ 1.8 µm (362 nm – 4.7 µm) and CMO^+^ CD63^+^ CD9^+^ 2.5 µm (362 nm – 6.2 µm). When considering the second phenotype: All CMO >MESF 50 1.4 µm (362 nm – 6.2 µm), CMO only 1.1 µm (362nm – 4.6 µm), CMO^+^ CD63^+^ CD9^−^ 1.7 µm (362 nm – 4.5 µm), CMO^+^ CD63^−^ CD9^+^ 1.6 µm (362 nm – 5.6 µm) and CMO^+^ CD63^+^ CD9^+^ 2.2 µm (362 nm – 6.1 µm). In each case the median diameter increases in size with accumulating EV marker expression.

When considering all 5 replicates; we compared the mean diameter for each subgroup (Figure 6 C iii). For the first group (left panel): CMO only 0.89 µm (+/− 0.03 SEM), CMO^+^ CD63^+^ CD9^−^ 1.53 µm (+/− 0.1 SEM), CMO^+^ CD63^−^ CD9^+^ 1.76 µm (+/− 0.17 SEM) and CMO^+^ CD63^+^ CD9^+^ 2.19 µm (+/− 0.18 SEM). Those EVs expressing either CD63 (*p* < 0.0004) or CD9 (*p* < 0.0010) were significantly larger than EV which didn’t (CMO only). For the second subgroup (right panel) the mean diameters were as follows: CMO only 0.84 µm (+/− 0.04 SEM), CMO^+^ CD63^+^ CD9^−^ 1.06 µm (+/− 0.12 SEM), CMO^+^ CD63^−^ CD9^+^ 1.31 µm (+/− 0.12 SEM) and CMO^+^ CD63^+^ CD9^+^ 2.02 µm (+/− 0.07 SEM). In this group, those EVs expressing CD63 only were not significantly larger than those which did not. However, EVs expressing LAMP1 were larger (*p* < 0.005). In both cases, those CMO ^+^ EVs co-expressing 2 EVs markers: either CD63 and CD9, or CD63 and LAMP1 (blue) were significantly larger than those which do not carry the markers screened in this study. (**** *p* < 0.0001. Un-paired t-test).

## DISCUSSION

Our principal aim was to develop a standardised method for the isolation and characterisation of individual large EVs, which could be further developed for phenotyping large EVs from clinical samples. The value of the Imagestream to the field of EV characterisation has been explored elsewhere (14, 15, 19). However, reports focussing on the large EV populations; frequently discarded during small EV isolation protocols, are rarely present in the current literature.

By using *in vitro* cultures, we were able to use the ideal conditions to generate EVs and optimise experiments. Specifically, by culturing in serum free media, we eliminated contamination from bovine EVs present in FBS (16) and subsequent false positive fluorescence signals from serum lipoproteins (17). Nevertheless, harvesting large EVs from any source presents challenges as cells or cell debris, including intracellular vesicles released due to parent cell membrane rupture from early centrifugation steps, can contaminate the subsequent EV pellets. For the EVs to be truly extracellular prior to isolation, the outer membrane of accompanying cells must not be ruptured by mechanical or chemical means during initial harvest. The centrifugation speeds we have used here are low compared to some commonly reported EV isolation protocols (20) to specifically preserve large EV membrane integrity. Electron microscopy remains the sole technique that can examine individual EVs and EV preparations for sheared cell fragments, but it is neither quantitative nor high throughput. It is necessary therefore to use a combination of techniques to explore the quantity, quality and biology EVs. All techniques, many originally designed for analysing cells, have technical challenges when applied to considerably smaller entities. For flow cytometry, background scatter events due to particles in the sheath fluid are an anticipated phenomenon which is rarely reported. In the work we report here, a high level of background appeared within the same gate as unstained EVs and persisted despite 0.2 µm filtration of sheath fluid. Our protocol is therefore reliant on strong and uniform, membrane-bound fluorescence labelling in order to assign an initial gate that separates potential EVs from speed beads or background scatter events. However, we found that the lower fluorescence intensity threshold to define CMO positive events was not clearly distinct from the instrument background. This was likely due to a combination of EV size and relative fluorescence intensity. As recommended elsewhere (14) we used commercially available fluorescent beads of known intensity (Quantibrite) to provide a means to assign standardised units (MESF) and therefore a mathematical cut-off for our CMO+ gate. PE was the closest available fluorophore to CMO and used as a standard for channel 03 on the ISX. CMO is a membrane label incorporated into the plasma membrane, and emits a greater fluorescence compared with a target-specific, conjugated antibody. This hampered the use of FITC alongside CMO as the spectrally close fluorophores led to over compensation between channels 02 (FITC) and channel 03 (CMO). BV421 was a successful alternative but the difficulties encountered raised concerns about trying to further multiplex with additional fluorophores using this platform.

The challenges for choosing the correct technique(s) to analyse EVs have been well described elsewhere (21). A major challenge now is to adapt the protocol we describe for the analysis of large EVs from biological fluids. Clinical samples will be more complex: EVs from a single cell type are unlike clinical samples which contain EVs from numerous cells (10). Our SOP is likely to exclude most small EVs and exosomes, expected to be present in the supernatant discarded at the final step (2000 × g). Experiments to isolate these for comparison are on-going. One classification we examined in this study was the distinction from apoptotic bodies. Demonstration of intact EV membranes, a lack of fragmented nuclei staining (DAPI) and evidence that EVs were derived from viable cells supported our assertion that the EVs analysed were not apoptotic bodies or cell debris (22).

Our data show that large EVs are ubiquitous and whilst absolute quantification is not yet within reach, we demonstrated similar size profiles using two independent techniques: TRPS and ISX. We have previously identified EVs of up to 6 µm using immunofluorescence, ISX and TEM (11). However, validating the quantity and size range has only been possible using the qNANO instrument. The qNANO employs TRPS technology to quantify EV count in a given sample and assign a size relative to a calibration bead of known diameter. It is currently the only instrument which can provide this information across the large EV population which spans 250 nm up to 6 µm (from our cells). Other platforms are restricted to small EVs (<1 µm) due to the measurements being reliant on Brownian movement (e.g. Nanosight). We found the most prevalent EV populations to be around 250-450 nm; however we consistently detect EVs with a much larger diameter range. We remain cautious not to define these as oncosomes; as although derived from cancer cells, we have not yet demonstrated their oncogenic potential (4).

Whilst fluorescence intensity alone cannot be used to quantify protein expression levels due to low level antigen expression on EVs, we did observe patterns of differential expression. In EV literature, 3 principle markers are used to define EVs: CD63, CD9 and CD81. CD63 however has been identified as a pan-EV marker, present in all defined EV subgroups to date (6) and therefore CD63 was our preferred initial marker. However, large EVs are as yet poorly characterised and we found that CD63 was not the most abundant EV marker in our study. CD63 is a tetraspanin which has been used as a selection tool for immuno-capture experiments and also for tracking EV release (23, 24). Based on our observations, if a full EV repertoire was of interest, then a cocktail of multiple markers should be considered as screening with CD63 alone will likely fail to capture a significant proportion of EVs. We found LAMP1, previously identified on exosomes and EVs from a variety our cell types and biological fluids (25) was significantly more prevalent.

We also show here that the majority of large EVs did not express any of the three markers screened. We did observe a significant increase in the median diameter of individual EVs which expressed 1 or more markers compared to none (CMO only), in each of the experiments performed. Further, across all replicates, those EVs which co-expressed 2 markers (CD63 + CD9, or CD63 + LAMP1) were significantly larger than those without. Others have suggested that larger EVs are likely to accommodate a greater number of tumour-derived molecules than exosomes (26) and data presented here would support that hypothesis. Further investigations to define a broader panel of large EV markers followed by more comprehensive techniques such as proteomic profiling, would help to fully phenotype the large EVs population. From a clinical perspective, it is likely that large EVs will be a rich source of biomarkers of benefit to the study of human disease. Standardised protocols and instruments capable of measuring multiple markers are key to move the field forward and expand the interest from exosomes only.

We set out to develop an isolation protocol consisting of minimal manipulation and processing which may abrogate, mask or indeed elicit changes in EV structure or biology which could impact on any functional read outs in downstream experiments. The research we report here demonstrates that high resolution, high throughput imaging flow cytometry is an exceptional tool offering the unique ability to quantify and analyse individual events within heterogeneous EV populations.

